# Evolutionary trajectories of secondary replicons in multipartite genomes

**DOI:** 10.1101/2023.04.09.536151

**Authors:** Natalia O. Dranenko, Aleksandra D. Rodina, Yaroslav V. Demenchuk, Mikhail S. Gelfand, Olga O. Bochkareva

## Abstract

Most bacterial genomes have a single chromosome that may be supplemented by smaller, dispensable plasmids. However, approximately 10% of bacteria with completely sequenced genomes, mostly pathogens and plant symbionts, have more than one stable large replicon. Some secondary replicons are species-specific, carrying pathogenicity or symbiotic factors. Other replicons are common on at least the genus level, carry house-keeping genes, and may have a size of several million base pairs.

We analyzed the abundance and sizes of large secondary replicons in different groups of bacteria and identified two patterns in the evolution of multipartite genomes. In nine genera of four families, *Pseudoalteromonadaceae, Burkholderiaceae, Vibrionaceae*, and *Brucellaceae*, we observed a positive correlation between the sizes of the chromosome and the secondary replicon with the slope in the range of 0.6–1.2. This indicates that in these genera the replicons evolve in a coordinated manner, with comparable rates of gene gain/loss, hence supporting classification of such secondary replicons as ‘chromids’. The second, more common pattern, features gene gains and losses mainly occurring in the primary replicon, yielding a stable size of the secondary replicon. Such secondary replicons are usually present in only a low fraction of the genus’ species. Hence, such replicons behave as ‘megaplasmids’. A mixed situation was observed in symbiotic genera from the *Rhizobiaceae* family where the large secondary replicons are of stable size, but present in all species. These results may provide a general framework for understanding the evolution of genome complexity in prokaryotes.

**Significance:** Large secondary replicons are observed in representatives of many taxonomic groups of bacteria. Traditionally, they are referred to as *second chromosomes, chromids*, or *megaplasmids*, with little consistency, in particular because their evolution remains understudied. Here we demonstrate that the sizes of secondary replicons follow two main evolutionary trends: replicons whose size scales linearly with the size of the main chromosome (the suggested term *chromids*) typically contain numerous essential genes (rRNA, tRNA, ribosomal proteins), while large secondary replicons of stable size (termed *megaplasmids*) contain fewer or none such genes.

## Introduction

Classical microbiology assumed bacterial genomes to be a single circular chromosome possibly supplemented by small nonessential circular plasmids. Since 1979, when linear plasmids in *Streptomyces* were discovered (Hayakawa et al. 1979), these assumptions started to change. Ten years later two chromosomes in the *Rhodobacter* genome were described (Suwanto and Kaplan 1989). This type of genome organization was named multipartite genomes, with about 10% of sequenced bacterial species being of that type (diCenzo and Finan 2017).

When a bacterial genome is formed by several large elements, the largest one, that holds the main fraction of house-keeping genes, is considered the primary chromosome, other parts being classified as chromids, megaplasmids, or plasmids based on their size and gene content (for details, see (diCenzo and Finan 2017)), but in the absence of universal definitions this classification is mainly based on arbitrary thresholds (Burton et al. 2013; Weiser et al. 2019; Cazares et al. 2020; Bottacini et al. 2015; diCenzo and Finan 2017). The term *chromid* was introduced by (Harrison et al. 2010) to describe large secondary replicons replicated by the plasmid-type mechanism but containing essential genes found in chromosomes in other species. Chromids were observed mostly in *Proteobacteria* but also in some distant genera, such as *Prevotella, Leptospira*, and *Deinococcus* (Harrison et al. 2010). The term *megaplasmid* is applied to plasmids larger than ∼100 kb (Burton et al. 2013; Weiser et al. 2019; Cazares et al. 2020; Bottacini et al. 2015; diCenzo and Finan 2017). Genomes of some species may hold several secondary replicons of different types, *e*.*g Burkholderia cepacia* that has a chromid and a huge essential megaplasmid (Bochkareva et al. 2018).

Evolutionary advantages of the multi-chromosomal genome organization are not clear. One hypothesis suggests that it has evolved to allow for gene accumulation and genome size expansion with two opposite directions of selection, one driving accumulation of new genes to increase the organism fitness, and the other reducing the genome to increase the replication rate. Multiple replicons may replicate simultaneously allowing for faster bacterial cell division even for large genomes (Guo et al. 2003), although there is no strong correlation between the number of replicons and the replication time (diCenzo and Finan 2017). The bacterial genome size varies from 112 kb for obligate symbionts (Bennett and Moran 2013) to ∼15 Mb for some free-living organisms (Han et al. 2013). While multipartite genomes are larger on average, large single-chromosome genomes exist, and less than one third of genomes larger than 6Mb are multipartite (diCenzo and Finan 2017). Therefore, the genome size is not uniquely driven by the number of replicons. Another hypothesis suggests that secondary replicons are expected to be under weaker selective pressure and have more hotspots of incorporation of new horizontally transferred genes (diCenzo et al. 2019). At that, secondary replicons may act as “test beds” that accumulate new genes and allow them to evolve more rapidly.

Here we study the relationships between the replicons’ sizes in multipartite genomes and interpret these observations in the context of evolution.

## Results

### Genome sizes

In the absence of a universally accepted definition of a multipartite genome, we define ‘multipartite genome’ as a genome with at least one secondary replicon longer than 500 Kbp, the threshold set at the first local minimum of the distribution of the secondary-replicon sizes (Supplementary Figure 1). We analyzed all representative bacterial genomes from RefSeq to select genera with at least one reference multipartite genome. Then we filtered out genera that comprised less than six genomes (see Methods). Thus, we analyzed genome configuration and size for 547 species from 36 genera.

The smallest genome (2.5 Mb) was of *Prevotella* sp. F0039 and the largest genome (12.2 Mb) was of *Streptomyces sp*. RLB1-9 (Supplementary Table 1). In agreement to previous observations, we did not observe any association between the genome size and taxonomy (Figure 1).

**Figure 1.**
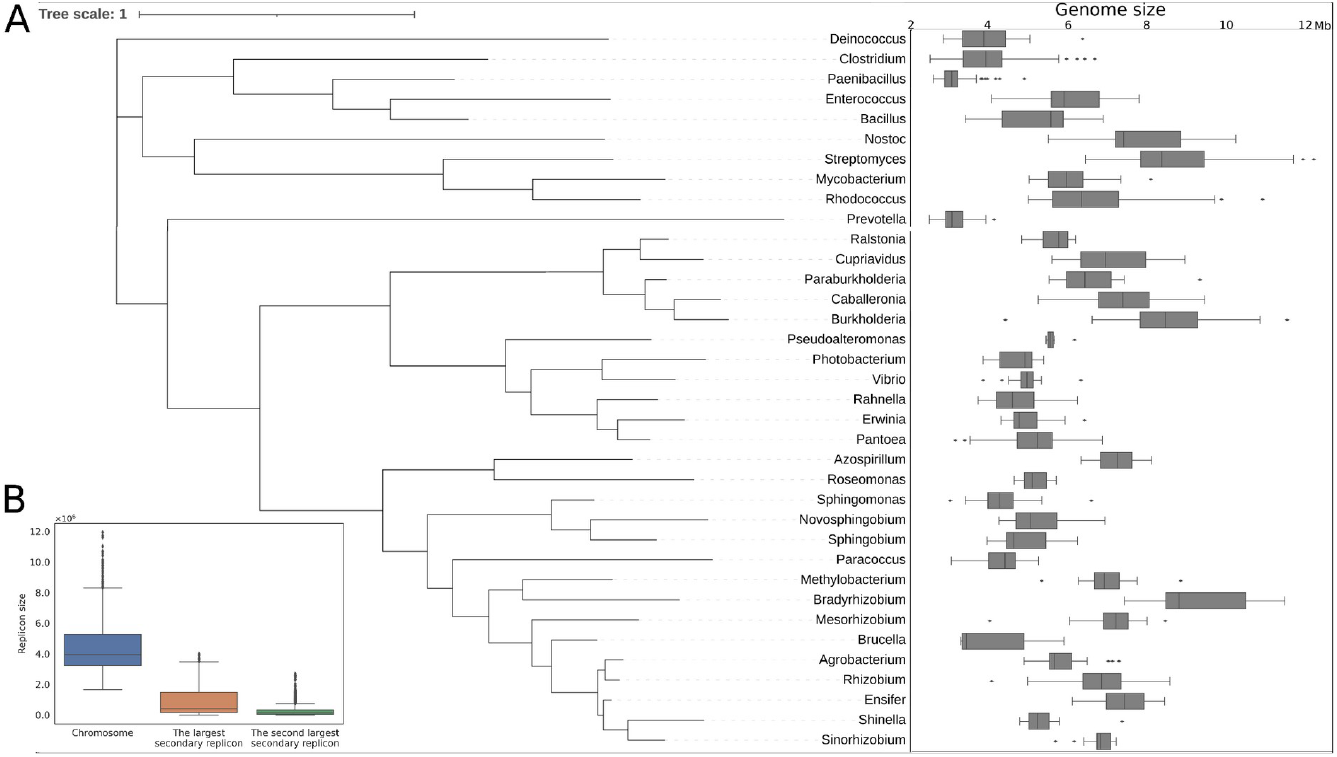
Replicon sizes in 36 genera with multipartite genomes. (A) Genome sizes of strains from genera that have members with multipartite genomes (see Methods). The cladogram topology follows the NCBI taxonomy (Schoch et al. 2020). (B) Replicon sizes; blue shows chromosomes; orange, the largest secondary replicon; green, the second-largest recondary replicon.

**Table 1.**
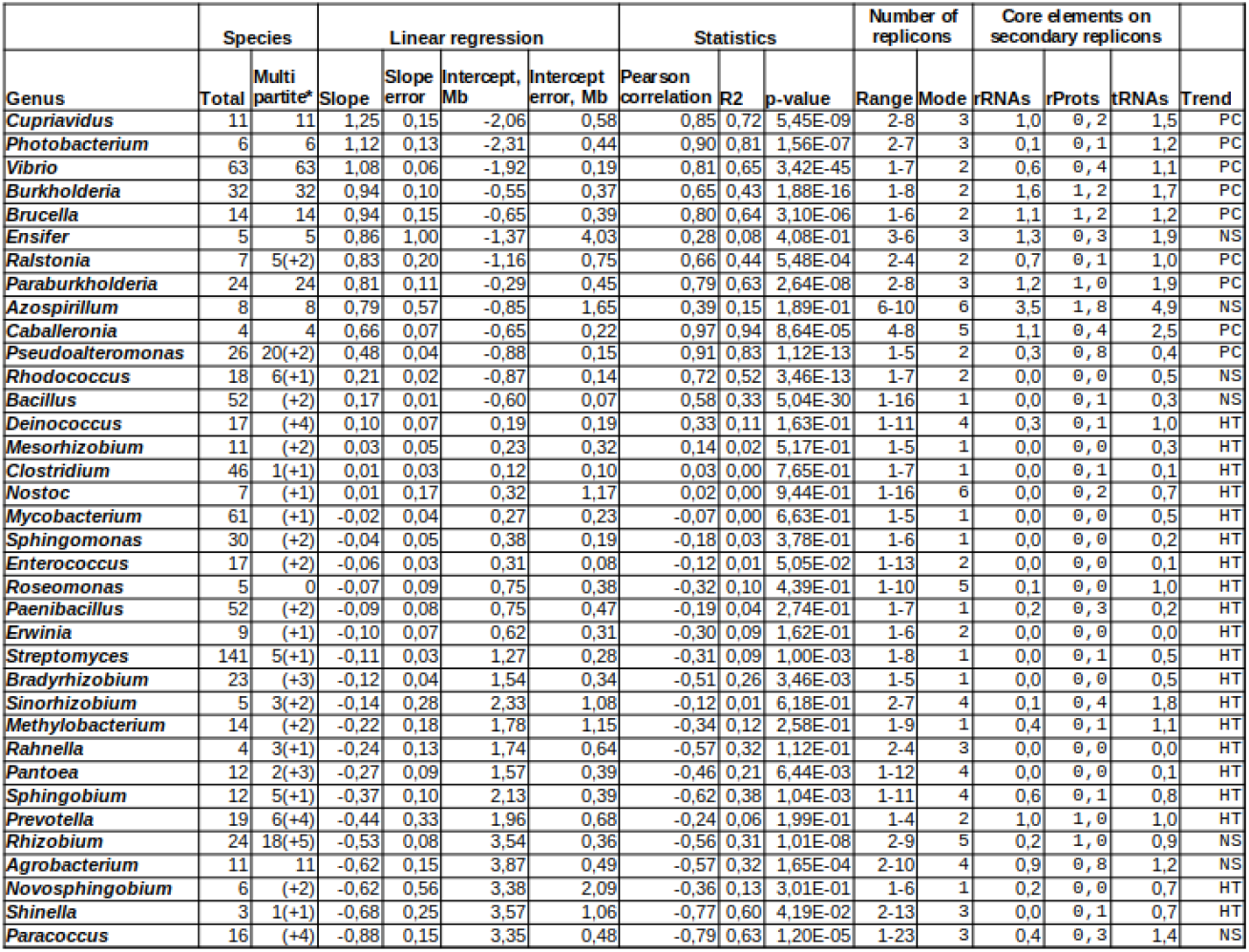
Abundance of multipartite genomes and the coefficients of linear regression between the sizes of the chromosome and the largest secondary replicon, per genus. Columns: Total — the number of species in the genus; multipartite — the number of species with strains having multipartite genomes (for species containing less than 5 strains the value is given in parentheses); slope and intercept — coefficients of the linear regression; slope_error and intercept_error — standard error of the estimated slope and intercept under the assumption of residual normality; Pearson correlation — the Pearson correlation coefficient; R2 — Coefficient of determination; p-value — the *p*-value for the null hypothesis that the slope is zero, using the Wald test with t-distribution of the test statistic; Range and Mode — the number of replicons, its range and mode value, in a genus; rRNAs, rProts, and tRNAs — the mean number of secondary replicons with the respective features, in a genus; Trend — strong positive correlation (PC), approximately constant size of the secondary replicon (HT), or non-significant positive correlation or non-homogeneous genera (NS).

### Replicon sizes

We compare the sizes of chromosomes and secondary replicons in genera with multipartite genomes. The chromosomes in some genera reach 12 Mbp, and the secondary replicons may be of length 4 Mbp (Supplementary Figure 1,2). The intervals of sizes of the chromosome and the largest secondary replicon is very large, while the second-largest secondary replicons rarely reach the chromosome size.

### Relationships between replicon sizes

Visual comparison of sizes of the chromosomes and the largest secondary replicons revealed two trends: the *horizontal* trend formed by plasmids with a stable size and the *diagonal* trend reflecting the consistent growth or reduction of both replicons (Figure 2).

**Figure 2.**
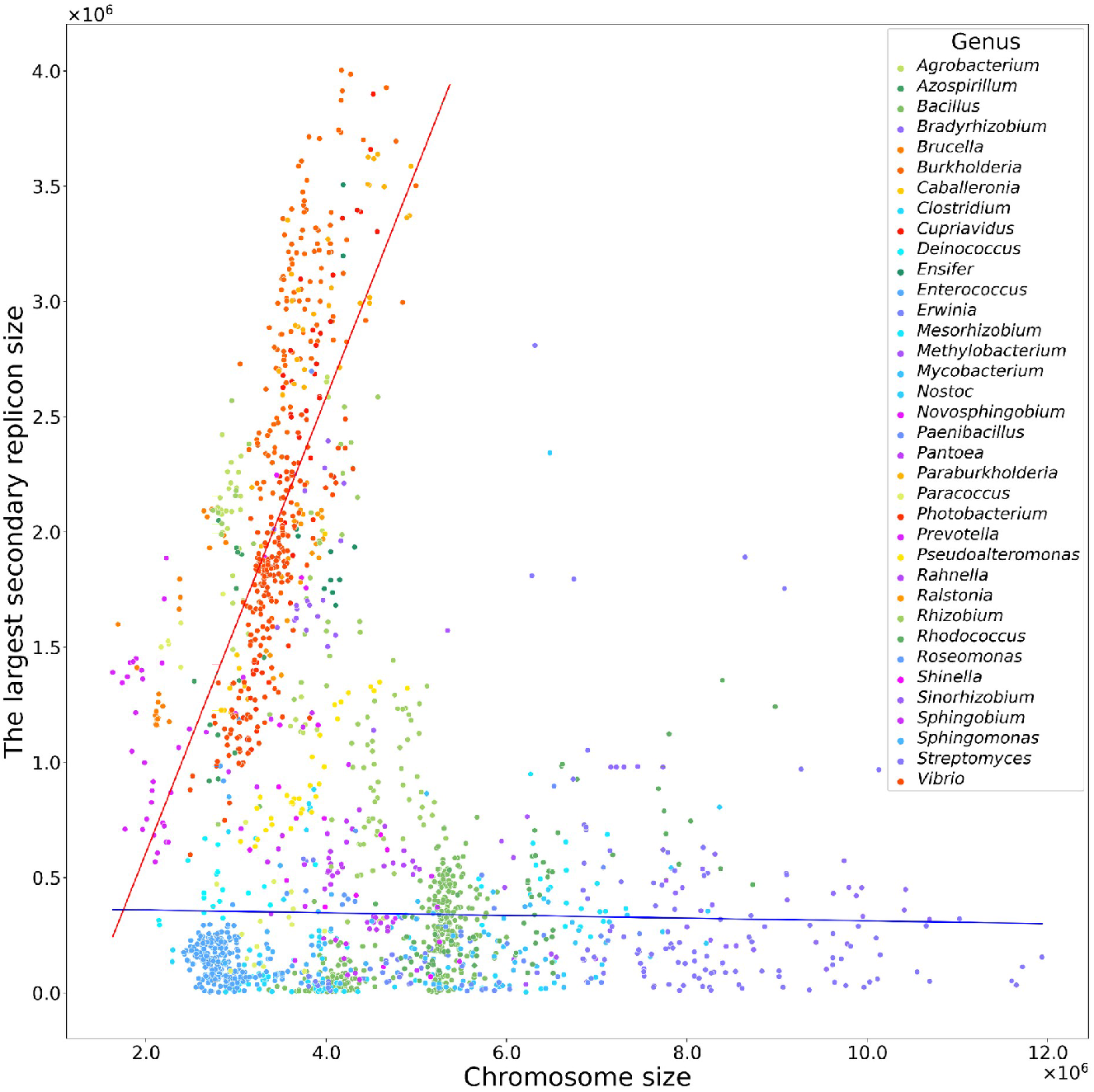
The chromosome size (horizontal axis) and the size of the largest secondary replicon (vertical axis) for strains with multipartite genomes from 36 genera. The dot color shows the bacterial genus (see the legend). Blue-purple colors show genera with significant positive correlation, yellow-red colors show genera with approximately constant size of the secondary replicon, green colors show genera with non-significant positive correlation or non-homogeneous genera.

To confirm this hypothesis, we performed linear regression analysis for each genus separately (Supplementary Figure 3, Table 1). We observed a significant positive correlation between the sizes of chromosomes and the largest secondary replicons in the genera where the multipartite genome organization is common for all or at least most species (Figure 3A,B). In turn, in genera where species with multipartite genomes are rare, we observed no correlation (Figure 3C).

**Figure 3.**
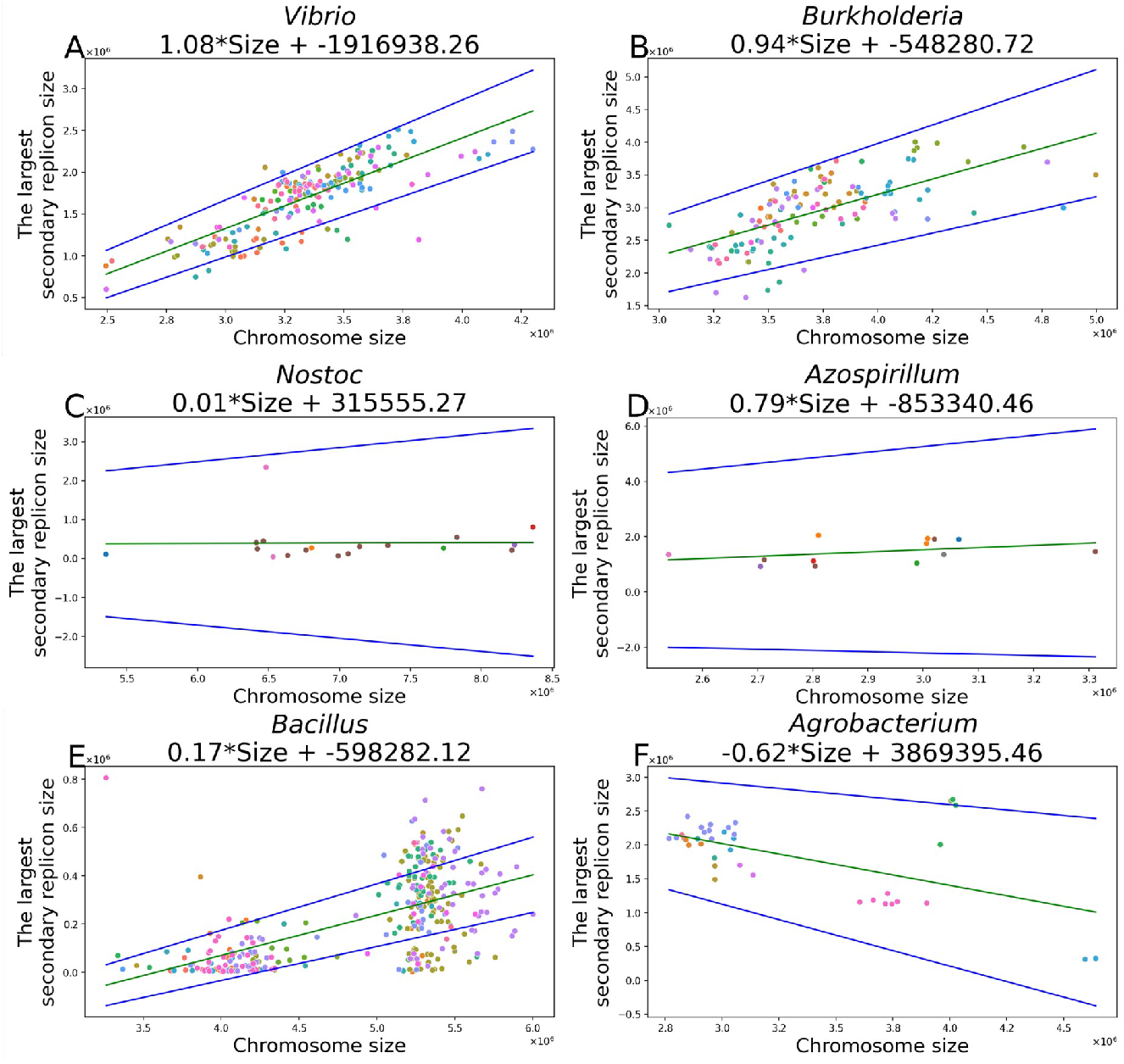
Linear approximation of the relationships between sizes of the chromosome and (C) *Nostoc*, genus with an approximately constant size of the secondary replicon (slope regression has a high variance; *r* = 0.4, slope coefficient 0.79 ± 1.14). (E,F) *Bacillus* and *Burkholderia* (*r* = 0.8, slope coefficient 0.94 ± 0.2), genera with a strong positive correlation; coefficient 0.01 ± 0.34); (D) *Azospirillum*, genus with a not-significant correlation (linear *Agrobacterium*, genera formed by several clusters of strains (the linear regression does not make sense). Green line reflects the resulting slope coefficient, blue lines reflect the 95% confidence interval for the slope.

However, there were several exceptions. *Ensifer and Azospirillum*, genera with the multipartite genome organization being common for all species, demonstrated positive but non-significant correlation (Figure 3D). A more interesting case was that of heterogeneous genera with significant positive or negative correlation (*Paracoccus, Agrobacterium, Rhizobium, Bacillus, Rhodococcus*). Linear regression is not applicable in this case, as these genera are formed by diverse subgroups of strains yielding separate clusters in the scatterplots (Figure 3E,F). Indeed, it is well known that *Bacillus* spp. are highly diverse and should be reclassified into several genera (Gupta et al. 2020). These genera were excluded from further analysis. the largest secondary replicon in: (A, B) *Vibrio* (*r* = 0.65, slope coefficient 1.08 ± 0.12) and

This procedure allowed us to classify the genera into two groups:

- significant positive correlation was detected in *Pseudoalteromonas, Burkholderia, Caballeronia, Cupriavidus, Paraburkholderia, Ralstonia, Photobacterium, Vibrio, Brucella*.
- non-significant or no correlation was observed in *Bradyrhizobium, Clostridium, Deinococcus, Enterococcus, Erwinia, Mesorhizobium, Methylobacterium, Mycobacterium, Nostoc, Novosphingobium, Paenibacillus, Pantoea, Prevotella, Sinorhizobium, Rahnella, Roseomonas, Shinella, Sphingobium, Sphingomonas, Streptomyces*.

Separate linear regression analysis for the groups (Figure 2) yielded two trends. For the first group *S*=0.99×*C*–1373734 (*C* is the chromosome size, S is the size of the largest secondary replicon), that is, *S*≈*C*+1,4×10^6^, and the correlation *R*=0.63. For the second group, *S*=– 0.006×*C*+370674 with *R*=–0.03 and the 95% confidence interval for the correlation being (– 0.0193, 0.0076); so this trend is essentially a horizontal one.

## Resulting trends

To get a general view, for each genus we constructed a Gaussian function with the mean equal to the slope coefficient and the standard deviation equal to the standard deviation of this coefficient (Figure 4A). Then we built a sum of these Gaussian functions (Figure 4B). The resulting distribution was bimodal, with the first peak at –0.05 and the second peak at ≈1.0. Hence, indeed, secondary replicons can be classified into two groups based on the relationships between their size and the chromosome size, with *megaplasmids* having no correlation, and *chromids* featuring a strong positive correlation.

**Figure 4.**
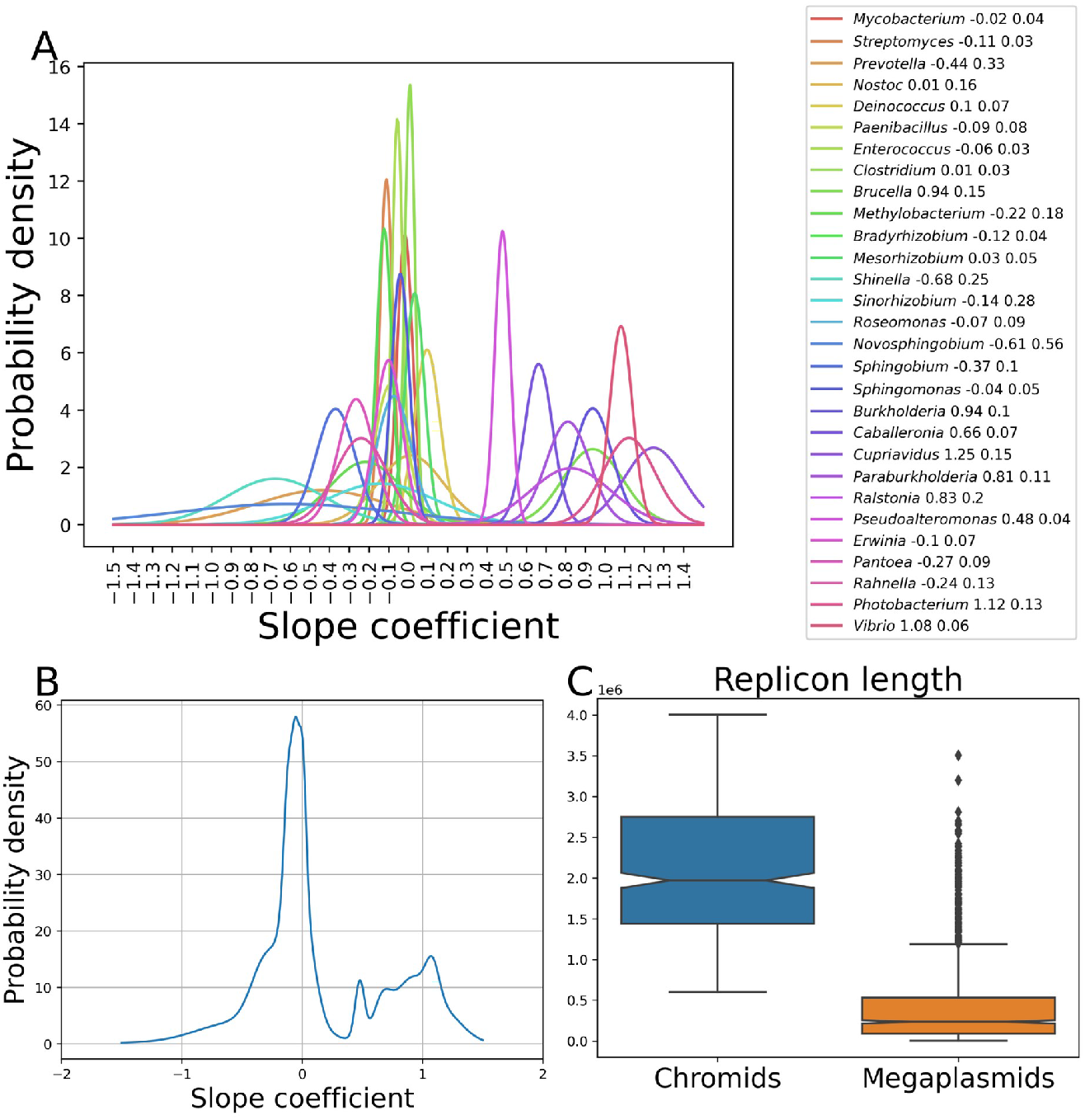
Summary of the regression analysis. (A) Gaussian functions derived from the slope coefficients (see the text), each colored line corresponds to a genus (listed in the legend; the numbers are the mean and the standard deviation). (B) Sum of the Gaussian functions. (C) Size distribution of the replicons classified as chromids and (mega)plasmids according to the regression analysis.

Note that replicons classified as chromids have sizes in the range 0.5-4 Mb, while megaplasmids do not exceed 3.5Mb and many are shorter than 500 kb (Figure 4C). Thus, the selection of genera with at least one 500 kb secondary replicon indeed has been an appropriate initial filter to study the organization of multipartite genomes.

### Contribution of other replicons

If gene distribution between the components of a genome is shaped by selection forces, the total number of replicons in a genome may affect the relationships between their sizes. It may explain the absence of correlation in *Rhizobia* as they possess many large replicons in comparison to *Vibrio*, typically having only two large replicons.

In all genera with the largest secondary replicons characterized above as ‘megaplasmids’, including *Rhizobiaceae* family, no correlation between the chromosome size and the total size of secondary replicons was observed (Supplementary Figure 4). Moreover, some genomes with chromids contain additional replicons of size comparable to that of chromids; their sizes do not demonstrate any consistent behavior (Supplementary Figure 5). In particular, in addition to a chromosome and a chromid, species of the *Burkholderia cepacia* complex (*Bcc*) **possess a common megaplasmid (∼1 Mb) involved in virulence and stress tolerance** (Agnoli et al. 2012). No significant difference between trends in *Bcc* with those of other *Burkholderia* was observed, demonstrating that acquisition of megaplasmids had not affected the relationship between the chromosome and chromid sizes (Supplementary Figure 6).

### rRNA, tRNA, and ribosomal protein genes

We compared the occurrence of three categories of *core* genes, namely rRNA genes, tRNA genes, and genes encoding ribosomal proteins (r-protein genes) in chromids and megaplasmids. As the chromids are bigger than 0.5 Mb while the megaplasmids are usually smaller (Figure 5A), we avoid biases by dividing secondary replicons into three groups based on their size and type: chromids (487 replicons), megaplasmids longer than 500 kb (385 replicons), and megaplasmids shorter than 500 kb (1025 replicons).

**Figure 5.**
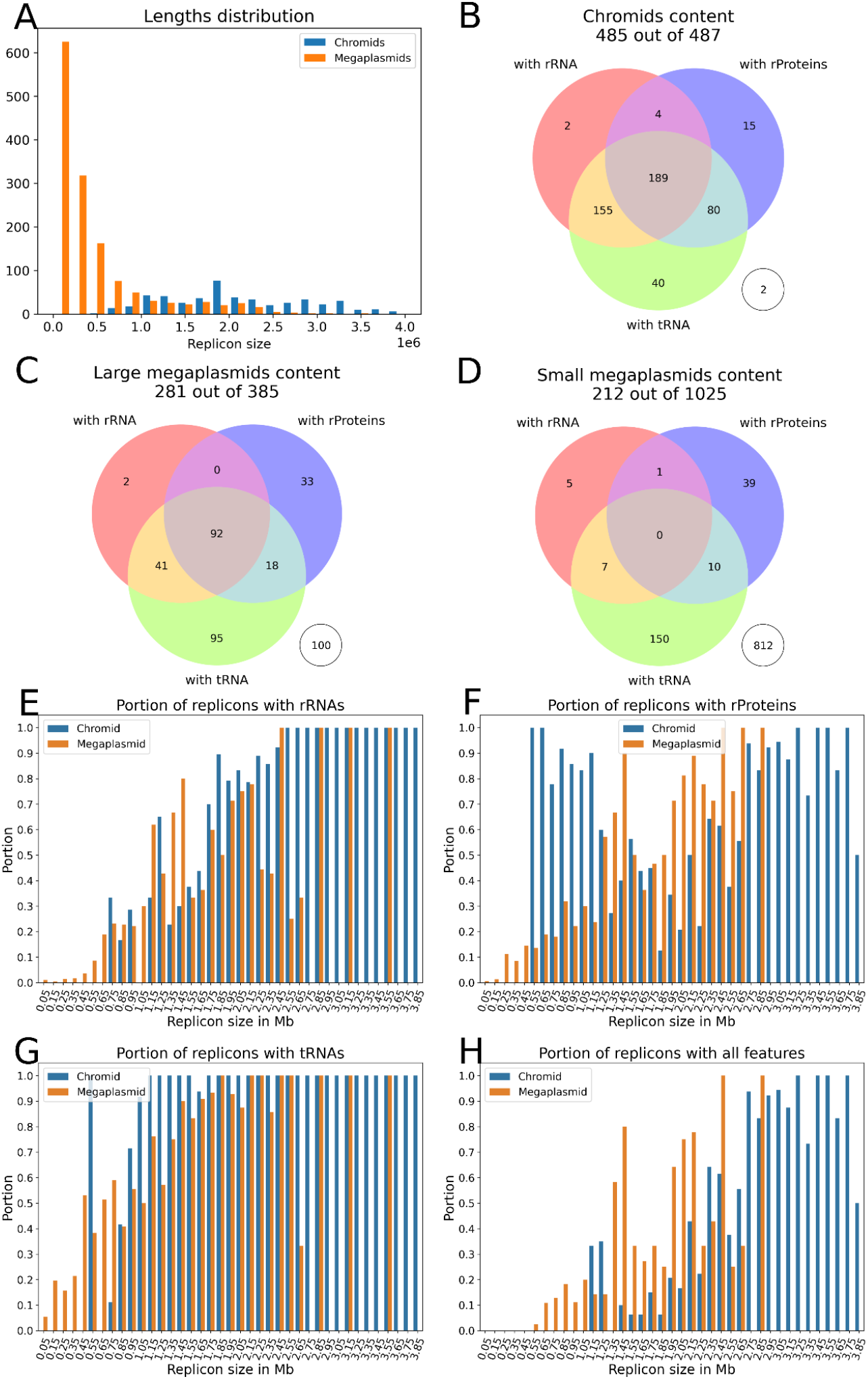
Occurrence of core features in secondary replicons. (A) Length distribution of secondary replicons characterized as chromids and megaplasmids based on the regression analysis. (B–D) Venn diagrams showing the occurrence of rRNA, tRNA, and r-protein genes encoding ribosomal proteins in secondary replicons characterized as chromids (B) and large (C) and small (D) megaplasmids; numbers in empty circles show the number of replicons with none of these genes. (E–G) Fractions of chromids and megaplasmids, depending on length, with rRNA genes (E), r-protein genes (F), and tRNA genes (G), at least one core gene of any category (H).

No such genes were found in only two chromids but more than one fourth of large megaplasmids (Figure 5B,C). In turn, ∼40% chromids (mainly from *Brucella, Burkholderia, Paraburkholderia*) and only ∼25% megaplasmids (mainly *Azospirillum* and *Prevotella*) featured at least one gene of each category. In small (mega)plasmids, we observed no replicons with at least one gene of each category, and only 10% of them possessed at least one of the core genes (Figure 5D).

Since the presence of the core genes could reflect the size of the secondary replicon rather than its type, we further calculated the fraction of replicons with core genes in cohorts defined by size (Figure 5E-H, Supplementary Figure 7). Indeed, the fraction of replicons with rRNA and tRNA genes is lower in smaller replicons. Surprisingly, although r-protein genes were rare in small megaplasmids, they were found in all size cohorts of chromids. In contrast to rRNA and tRNA genes, most r-protein genes are single-copy ones (Yutin et al. 2012) and their location in secondary replicons may explain the essentiality of the latter. To address this, we selected only those r-protein genes which were not present in chromosomes.

We observed such r-protein genes in the largest secondary replicons in *Brucella* (L33,S21), some *Vibrio* species (L20,L35), in of *Agrobacterium* species, and only several *Rhizobium* species (L36,L31,S1,S21,L28). In *Azospirillum*, the genus with an exceptionally high number of large replicons, the largest secondary replicon may contain (L33, L28) or (L13, S9) sets of r-protein genes.

The largest number of r-protein genes was observed in secondary replicons in the *Prevotella* genus. Note that genus *Prevotella* comprises species with a single ∼3Mb chromosome and species with two replicons, ∼2Mb and ∼1Mb in size. Moreover, in two-replicon species, rRNA, tRNA, and r-protein genes are proportionately distributed between the larger and smaller replicons indicating formation of such multipartite genome organization via chromosome division or translocation of a large fragment from the chromosome to the plasmid via intragenomic recombination.

## Discussion

The correlation between the chromosome and chromid sizes may reflect consistent growth or reduction of both replicons. In particular, in *Burkholderiaceae*, chromids show rapid accumulation of recently acquired genes associated with the adaptation to novel environments (diCenzo et al., 2019). Another example is an intracellular pathogen *B. mallei* that has relatively recently evolved from the extracellular pathogen *B. pseudomallei* (Bochkareva et al., 2018). This transition was accompanied by strong genome reduction, 12.5% in the chromosome and 31% in the chromid, that retained the proportion between the replicon sizes.

The second, more common trend demonstrates the absence of such correlation. This trend was mainly observed in genera with low fraction of multipartite genomes. Interestingly, it was also seen in megaplasmids of symbiotic genera from *Rhizobiaceae* (*Agrobacterium, Sinorhizobium, Rhizobium, Ensifer*, and *Shinella*), and *Azospirillaceae (Azospirillum)*. Note that large essential secondary replicons are common in *Rhizobiaceae* and *Azospirillum*, thus the absence of correlation could have been explained by the contribution of additional replicons. However, in these genera the correlation between the chromosome size and the total size of secondary replicons also was not observed. Moreover, acquisition of large megaplasmids by bacteria with chromids (e.g. *Bcc*) does not affect the chromosome-chromid size correlation.

The presented analysis may explain the origin and evolution of multipartite genomes. The size of a bacterial genome is associated with the lifestyle, with free-living bacteria having large genomes with open pangenomes, and intracellular pathogens having smaller genomes with smaller fractions of periphery (strain-specific) genes (Köstlbacher et al. 2021; Horesh et al. 2021). The correlation between the chromosome and the chromid sizes in *Pseudoalteromonas, Burkholderiaceae, Vibrionaceae*, and *Brucellaceae* both in genera and on the family level (Figure 6A-D) indicates that their chromids are involved in the gene gain/loss process, likely shaped by evolutionary selection under adaptation to different hosts and niches. In contrast, in *Rhizobiaceae* we observe no link between the chromosome and chromid sizes (Figure 6E). *Rhizobiaceae* are plant symbionts, characterized by the ability to modify plant development, with genes essential for nodulation and nitrogen-fixation located in megaplasmids and in chromosomal genomic islands (Yang et al. 2020). The absence of correlation may indicate specific functionality and, as a result, high conservation of composition of *Rhizobiaceae* secondary replicons.

**Figure 6.**
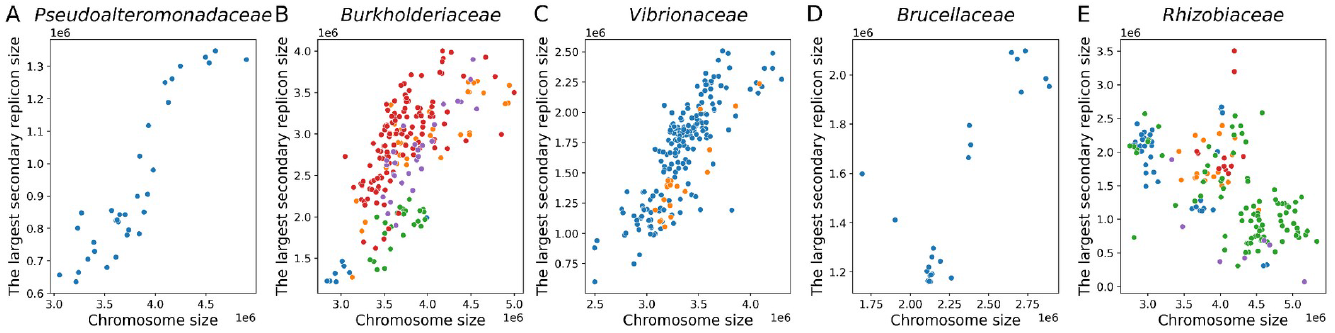
Relationships between the chromosome size and the largest secondary replicon size for five families with chromids: (A) *Pseudoalteromonadaceae* (*Pseudoalteromonas*), (B) *Burkholderiaceae* (*Caballeronia, Paraburkholderia, Ralstonia, Burkholderia, Cupriavidus*), (C) *Vibrionaceae* (*Vibrio, Photobacterium*), (D) *Brucellaceae* (*Brucella*), and (E) *Rhizobiaceae* (*Agrobacterium, Sinorhizobium, Rhizobium, Ensifer, Shinella*). Dot’s color reflects different genera.

Current classifications of secondary replicons of bacteria are rather arbitrary. The suggested criteria rely on the replicon size threshold and the presence of core (universal) or essential genes (Harrison et al. 2010; Hall et al. 2022). However, the definition of essentiality depends on the research question (e.g. with or without competition between strains) (Gerdes et al. 2003), whereas the core fraction of pan-genome depends on the taxonomy level of the latter. According to the strictest definition, chromids carry at least one gene that is essential for cell viability. This definition relies on experimental evidence that a replicon carries an indispensable core gene (diCenzo and Finan 2017). Here, we have considered three categories of core genes, common for all bacteria, tRNA and rRNA genes which are often found in multiple copies in many replicons (Santos and Del-Bem 2022; Gifford, Dasgupta, and Barrick 2021) and genes encoding ribosomal protein, usually located in the chromosome (Yutin et al. 2012). Indeed, the core genes were more frequent in replicons classified as chromids, compared to megaplasmids.

Our results suggest an evolutionary approach to the definition of chromids and megaplasmids based on general, consistent trends in their sizes. One caveat is that the performed analysis might be affected by several factors such as errors in genome assemblies and taxonomical misclassification. Moreover, replicons in multipartite genomes may be affected by recombination between *rrna* repeats resulting in large inter-replicon translocations (Treangen et al. 2009; Bochkareva et al. 2018) or replicon fusion (Sozhamannan and Waldminghaus 2020; Bochkareva et al. 2018; Sozhamannan and Waldminghaus 2020; Mori and Kanaly 2022). Thus, in order to further classify secondary replicons, the history of the gene flow between replicons should be considered in addition to the content and architecture of extant genomes.

## Materials and Methods

### Genomes and taxonomy

We started with all representative strains available in the NCBI RefSeq database (O’Leary et al. 2016). If at least one representative strain had a secondary replicon of size exceeding 0.5 Mb, we added the genus to the analysis (Supplementary Table 1).

### Dataset filtering

Thus, we analyzed the genome composition for 627 species from 84 genera. We reduced the dataset by selecting single representatives for groups of closely related strains. For this we clustered the genomes with respect to their replicon length (the differences less than 5%). Then we filtered out genera with less than 6 representatives. Finally, the dataset comprised 547 species from 36 genera (Supplementary Table 2).

### Analysis of outliers

Genome assemblies of several strains were significantly different from members of the same taxonomic group.

Secondary replicons in *Prevotella copri* DSM 18205 strain FDAARGOS_1573 and *Prevotella* sp. E2-28 were smaller than 210 kb, while other members of this species have only one replicon so these strains were considered as not multipartite.

In *Burkholderia cenocepacia* strain 895, *Burkholderia cenocepacia* strain VC12802, *Burkholderia vietnamiensis* strain AU1233, *Cupriavidus metallidurans* strain Ni-2, *Cupriavidus* sp. KK10, and *Brucella pseudogrignonensis* strain K8 we observed fusion of chromosomes and secondary replicons (Supplementary Figure 8). Here, we used Sibelia (Minkin et al. 2013) software to align and compare the outliers and the closest complete genome of the same genera. To identify the pair for each of the outliers, we build a distance matrix using Mash (Ondov et al. 2016).

The assemblies of *Pseudoalteromonas spongiae* UST010723-006, *Pseudoalteromonas piratica* strain OCN003, and *Pseudoalteromonas spongiae* strain SAO4-4 are multipartite but the sizes differ dramatically from other members of this genus that may indicate recent recombination events or assembly errors.

All these genomes were excluded from the analysis.

### Statistical analysis

Linear regression analysis was performed with the scipy.stats.linregress Python module. To construct the Gaussian functions for each genus, we used the slope coefficient as the mean μ and the slope standard error as the standard error σ in the Gaussian function

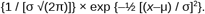

### Calculation of abundance of multipartite genomes in genera

We considered a genome as multipartite if it had a secondary replicon larger than 0.5 Mb. If more than 75% of strains forming a species had multipartite genomes, the species was marked as multipartite (Supplementary Table 3). Then, for each genus, we calculated the fraction of species with multipartite genomes (Supplementary Table 4).

## Supporting information

All supplementary figures

All supplementary tables

## Authors contribution

O.O.B. and M.S.G. conceived and designed the study. N.O.D., A.R., and Y. D. analyzed the data. N.O.D., O.O.B., and M.S.G. interpreted the results and wrote the graft. All authors read and approved the final version of the manuscript.

## Ethical approval

Not applicable.

## Consent for publication

Not applicable.

## Acknowledgements

The project was initiated at the Summer School of Molecular and Theoretical Biology (SMTB-2021), supported by the Zimin Foundation.

