## Supplementary material for "Evolutionary trajectories of secondary replicons in multipartite genomes": All supplementary figures

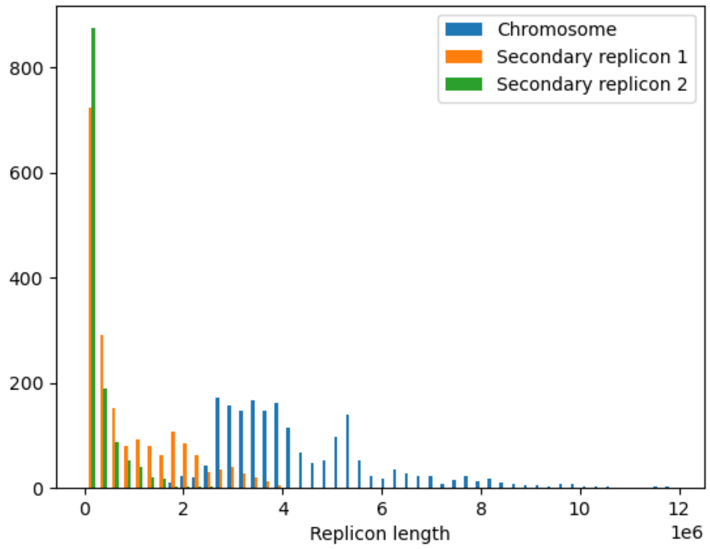


***Supplementary figure 1.*** Replicon sizes in the 36 genera with multipartite genomes. Blue shows chromosomes; orange, the largest secondary replicon; green, the second-largest recondary replicon.

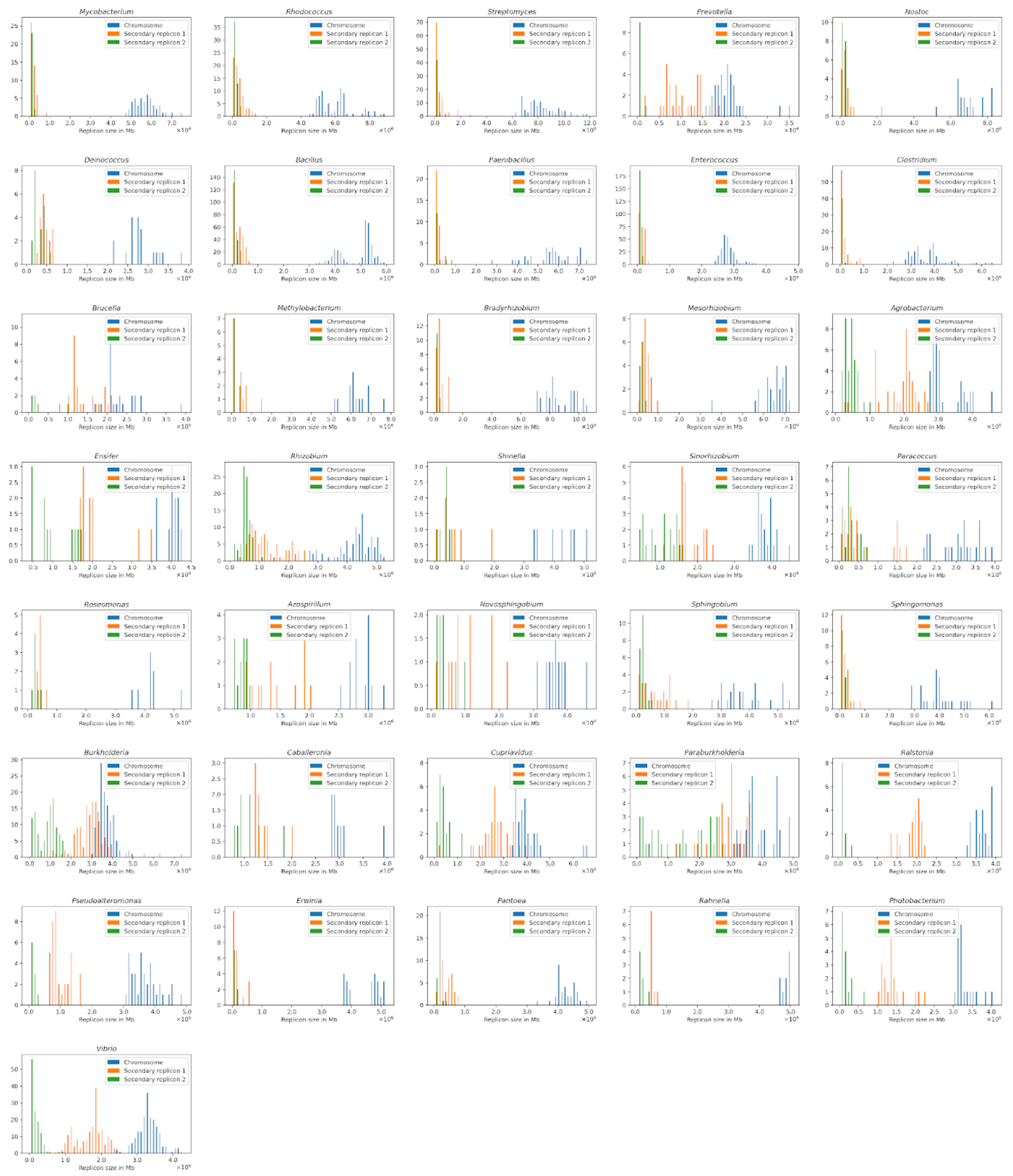


***Supplementary figure 2.*** Replicon sizes in the 36 genera separately. Notation as in Supplementary figure 1.

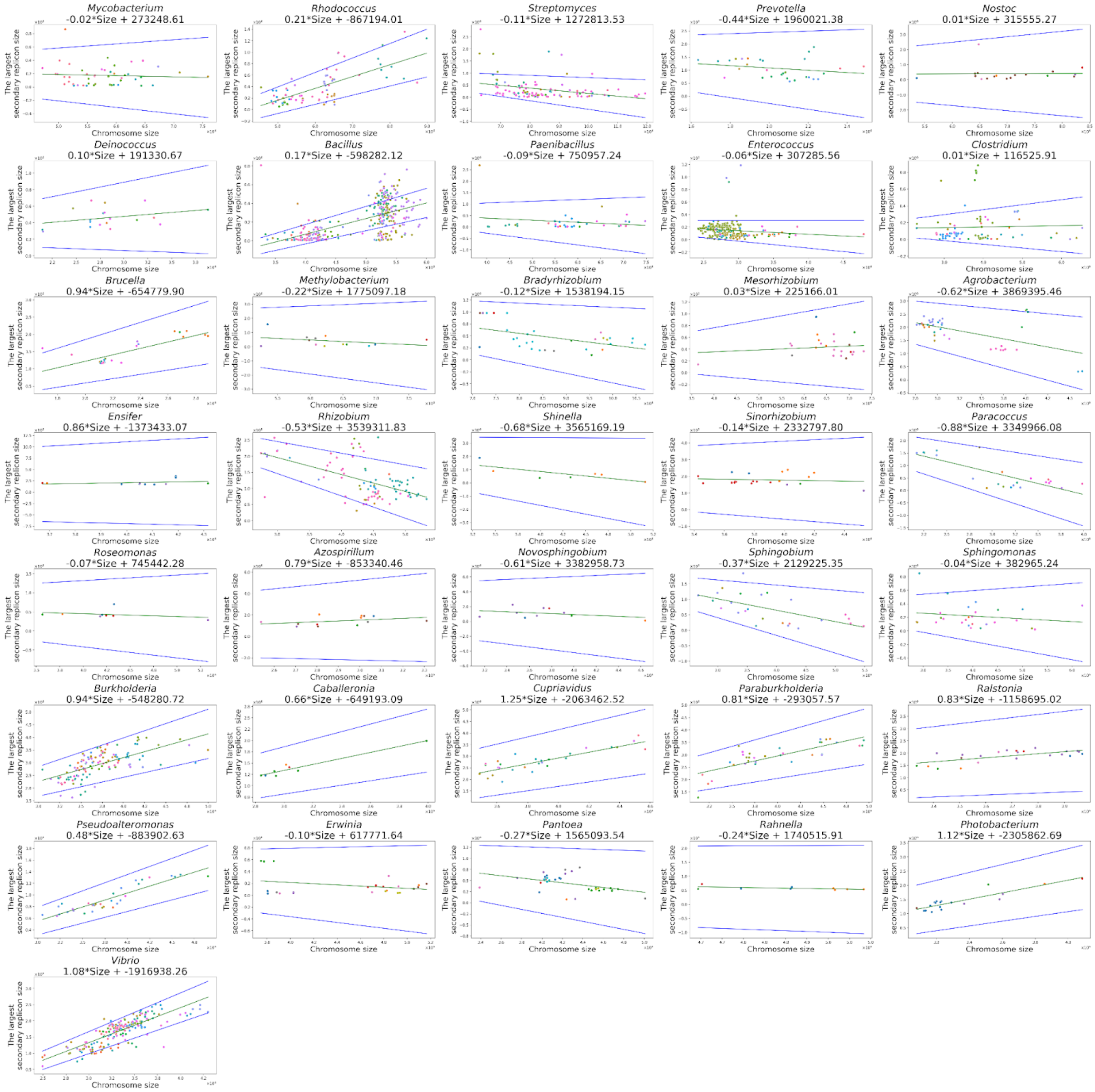


***Supplementary figure 3.*** Linear approximation of the relationships between sizes of the chromosome and the largest secondary replicon in the genera with multipartite genomes. Different species in a genus are marked by different colors. Green line reflects the resulting slope coefficient, blue lines reflect the 95% confidence interval for the slope. The slope coefficients and intercepts are shown in the titles.

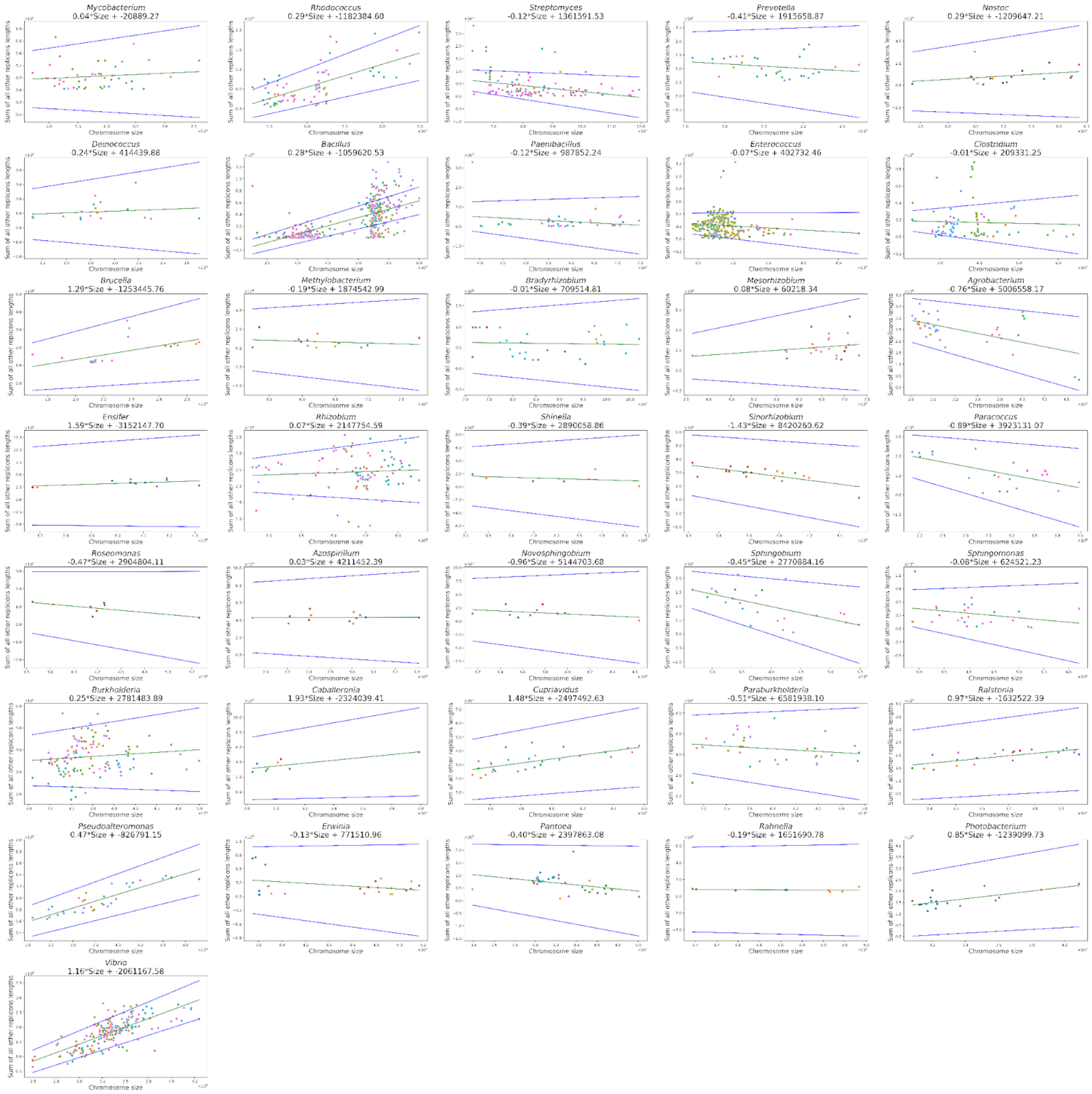


**Supplementary figure 4.** The chromosome size (horizontal axis) and sum of secondary replicon sizes (vertical axis).

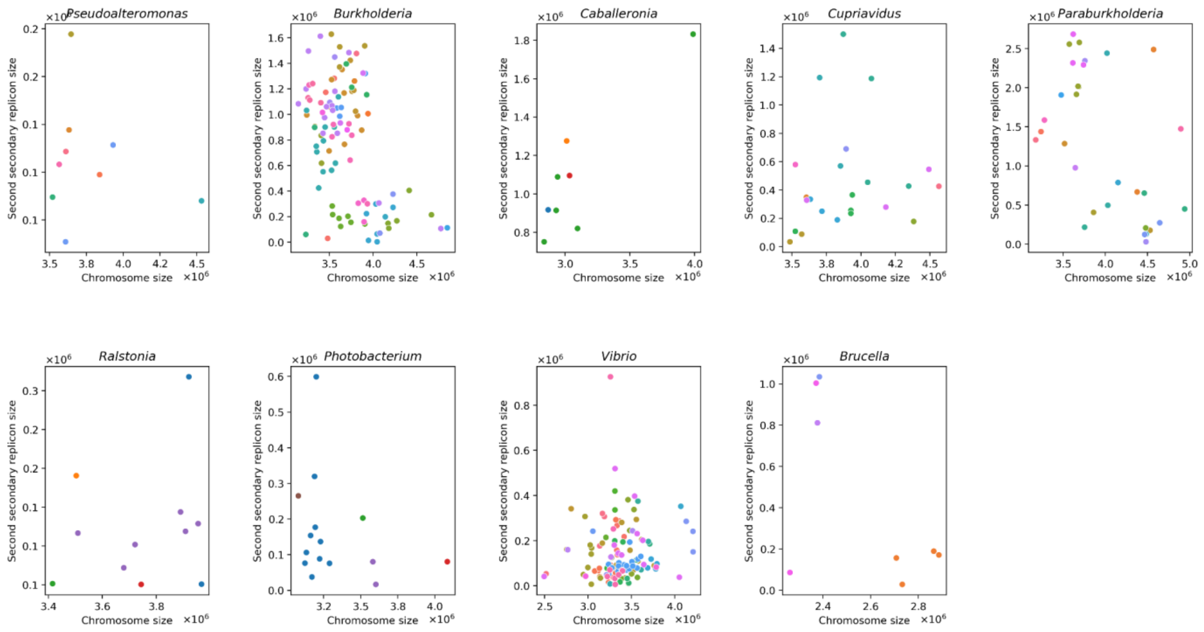


***Supplementary figure 5.*** Relationships between the chromosome and the second secondary replicon sizes for eight genera with the correlation between the chromosome and the largest secondary replicon was being observed. Different species in a genus are marked by colors.

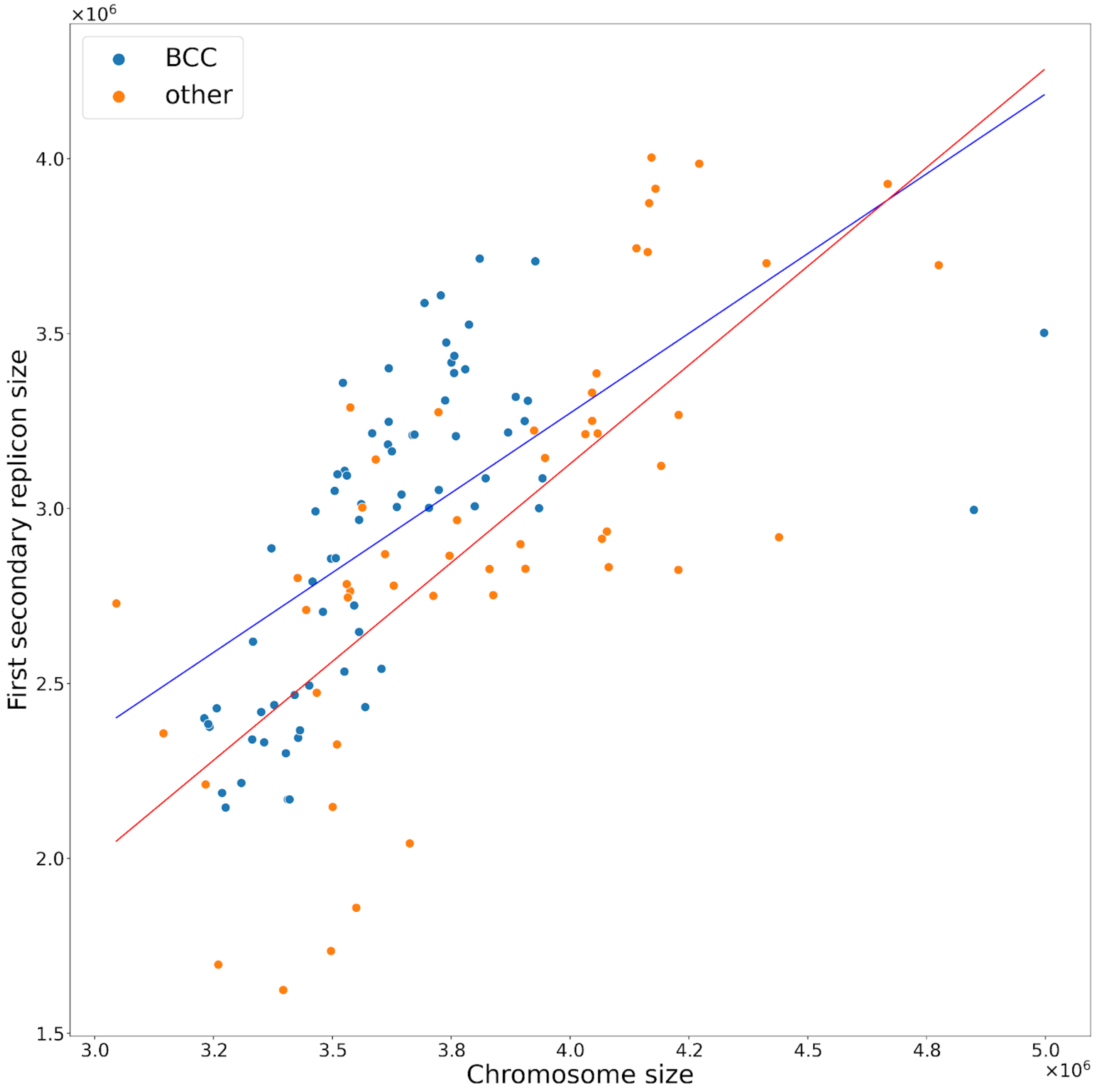


***Supplementary figure 6.*** The chromosome size (horizontal axis) and the chromid size (vertical axis) in *Burkholdeira* spp.; blue dots reflect *Bcc* strains with the large megaplasmid, red dots reflect other *Burkholderia*

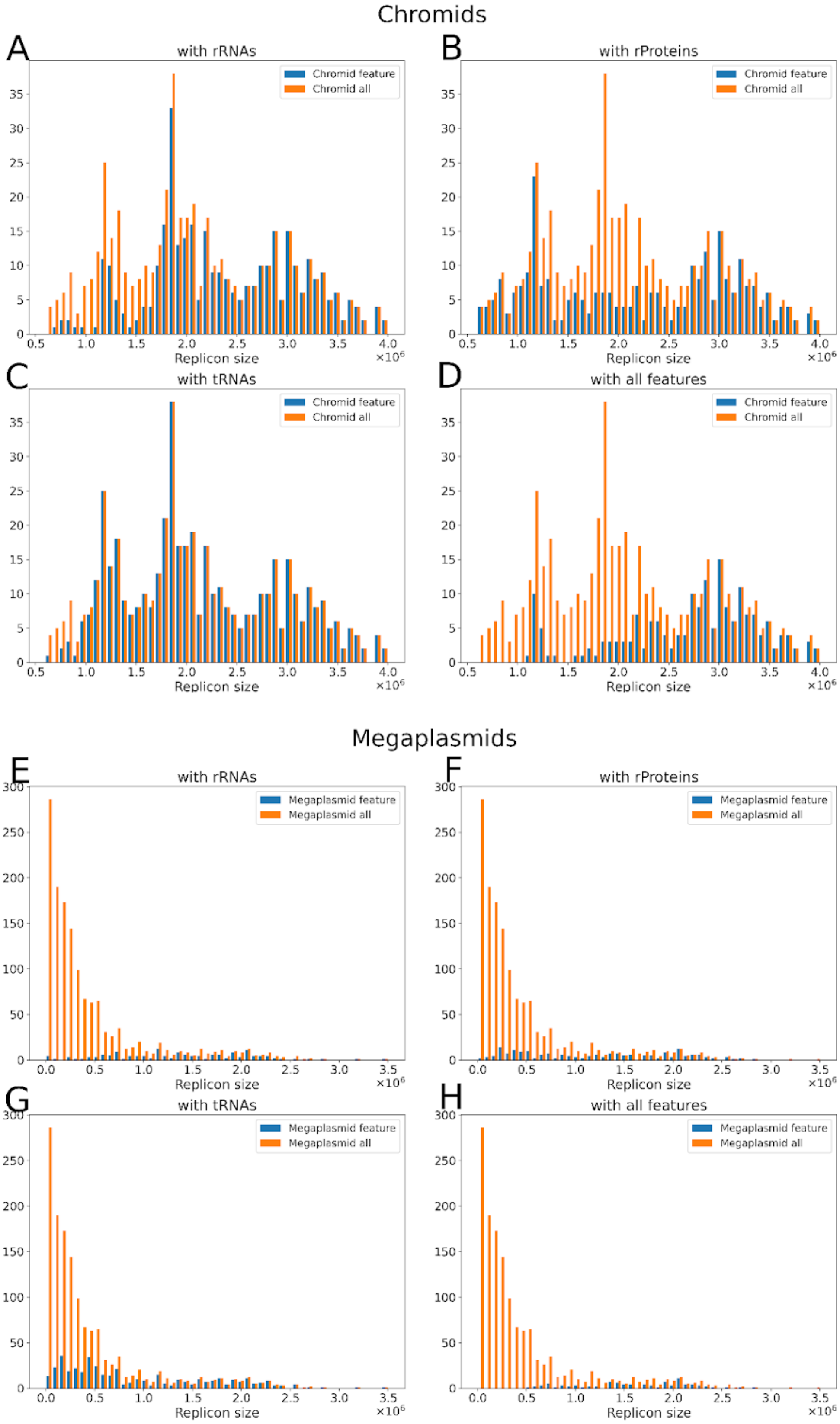

Chromids
A
with rRNAs
Chromid feature
Chromid all
Replicon size
B
with rProteins
C
with tRNAs
D
with all features
Megaplasmids
E
with rRNAs
Megaplasmid feature
Megaplasmid all
F
with rProteins
G
with tRNAs
H
with all features

**Supplementary figure 7.** Length distribution of the replicons with rRNAs, r-proteins genes, tRNAs, and at least one gene of each category in comparison to all replicons for chromids (A-D) and (mega)plasmids (E-H).

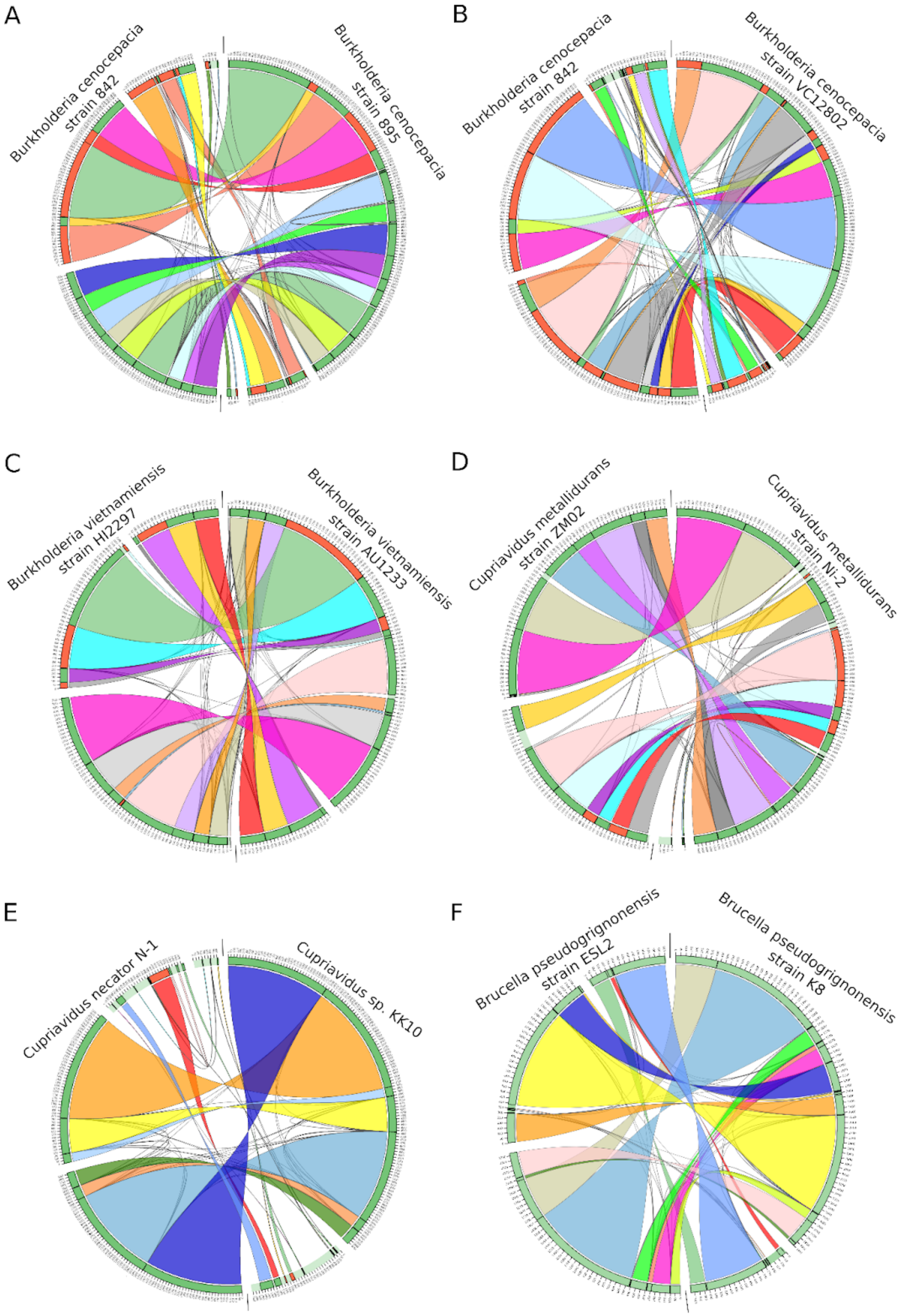

A
Burkholderia cenocepacia strain 842
Burkholderia cenocepacia strain 895
B
Burkholderia cenocepacia strain 842
Burkholderia cenocepacia strain VC12802
C
Burkholderia vietnamiensis strain HI2297
Burkholderia vietnamiensis strain AU1233
D
Cupriavidus metallidurans strain ZM02
Cupriavidus metallidurans strain Ni-2
E
Cupriavidus necator N-1
Cupriavidus sp. KK10
F
Brucella pseudogrignonensis strain ESL2
Brucella pseudogrignonensis strain K8

***Supplementary figure 8.*** Pairwise comparisons of strains using circos diagrams (A) *Burkholderia cenocepacia* strain 895 and *Burkholderia cenocepacia* strain 842; (B) *Burkholderia cenocepacia* strain VC12802 and *Burkholderia cenocepacia* strain 842; (C) *Burkholderia vietnamiensis* strain AU1233 and *Burkholderia vietnamiensis* strain HI2297; (D) *Cupriavidus metallidurans* strain Ni-2 and *Cupriavidus metallidurans* strain ZM02; (E) *Cupriavidus* sp. KK10 and *Cupriavidus necator* N-1.
